# Spatial Transcriptomics of Spinal Ependymoma in NF2-related Schwannomatosis

**DOI:** 10.1101/2024.03.03.583240

**Authors:** Noah Burket, Jia Wang, Hongyu Gao, Robert Bell, Chi Zhang, Yunlong Liu, Wade Clapp, Jignesh Tailor

## Abstract

Spinal ependymoma (SP-EPN) is a central nervous system (CNS) tumor that is associated with high morbidity. The only effective treatment for end-stage SP-EPN is surgery, but this is associated with high risk of injury to the sensorimotor spinal tracts and paralysis. There is a critical need to understand the cellular origins of this tumor so that disease models of tumor progression can be generated for drug development. Recent genomic studies with bulkRNA sequencing suggest the molecular signature of SP-EPN matches that of ependymal cells (EPCs). However, large-scale genomic studies can often misrepresent rare cancer stem cell populations within the tumor. In this study, we performed spatial transcriptomics (ST) on a SP-EPN resected from a patient with NF2-related neurofibromatosis to examine the spatial heterogeneity within the tumor. The SP-EPN sample exhibited cellular heterogeneity with diffuse expression of astrocytic and EPC markers, and smaller pockets with RGC or stem cell markers, as well as overlap between progenitor cell and mature cell markers. These findings suggest that there may be a developmental hierarchy within the EPC lineage in SP-EPN tumors, which may stem from aberrant radial glia cells.

## Introduction

Ependymomas are a heterogeneous group of tumors that can occur throughout the central nervous system (CNS).^1^ Spinal ependymoma (SP-EPN) is a distinct molecular entity that is found in the spinal cord and affects children and young adults. Previous studies suggest that *NF2* is a key genetic driver of spinal ependymoma (SP-EPN).^1–8^ Approximately, 50% of individuals with the inherited cancer predisposition syndrome NF2-related schwannomatosis (NrS) develop SP-EPN.^4,9^ Importantly, loss of heterozygosity of the *NF2* gene is seen in most spontaneous SP-EPN.^2,6,8^ Although SP-EPN are rare, occurring in approximately 1 in 250,000 individuals, they account for 20-30% of all tumors found in the spine and are difficult to treat.^10^ To date, there is no effective medical therapy for end-stage SP-EPN, and the surgical morbidity is high due to the proximity of these tumors to the spinal cord sensorimotor tracts and the risk of paralysis. Stopping the early lesions in patients with cancer predisposition such as NrS may be the ideal strategy to improve outcomes in this vulnerable group. This requires a better understanding of the cellular origins of SP-EPN and better disease models of tumor progression for drug testing.

In previous studies, ependymoma from different regions of the brain and spinal cord displayed molecular signatures that were like radial glia cells (RGCs) of the region from which they were derived.^11^ In addition, cancer stem cells isolated from ependymoma display a cellular phenotype reminiscent of RGCs, and these cells can be propagated in culture over long term.^11^ Further, these cells recapitulated the development of ependymoma when orthotopically transplanted in mice. These findings suggest RGCs may be candidate cells of origin of ependymoma.^11–13^ However, recent transcriptomic data suggest the expression profile of SP-EPN more closely matches that of mature ependymal cells (EPCs).^6^ In these experiments, the gene expression profile of SP-EPN through bulk RNA sequencing was aligned with the molecular signatures of cell clusters annotated from single-cell RNA sequencing of embryonic and adult spinal cord. Although the gene expression profile of the tumor resembled mature, adult spinal cord EPCs, bulk RNA sequencing data may not accurately represent rare populations of resident stem cells within the tumor, nor can it define the cellular hierarchy within the tumor.

In neural development, neuroepithelial cells (NECs) of the neural tube transition to RGCs, which provide a scaffold for early neurons to migrate and serve as precursors for other cell types in the spinal cord such as astrocytes and EPCs.^14,15^ We *hypothesized* a further developmental hierarchy exists in SP-EPN from RGCs to progenitors in the EPC lineage, with the majority terminating as mature, adult-stage EPC. To test this hypothesis, we studied the spatial heterogeneity of cells in SP-EPN from a patient with NrS using spatial transcriptomics (ST).

## Results

The sample was consistent with SP-EPN on histology. H&E staining showed a highly cellular, histologically homogenous, compact glial neoplasm (Fig. 1A, B). There was evidence of perivascular pseudorosettes on H&E and GFAP immunostaining (Fig. 1C), which is also typical of ependymoma. EMA staining demonstrated a microlumina in a low number of partially differentiated cells without true rosettes identified in this example (Fig. 1D). ST quality assessment showed good quality gene expression in most of our sample, and our DV200 was 27.5%. There was one cluster that showed low gene expression and reads (Fig. 2). This area was omitted from further examination. Unbiased cluster analysis of the spatially represented gene expression data revealed a highly heterogenous tumor with molecularly distinct areas (Fig.3).

**Fig. 1.**
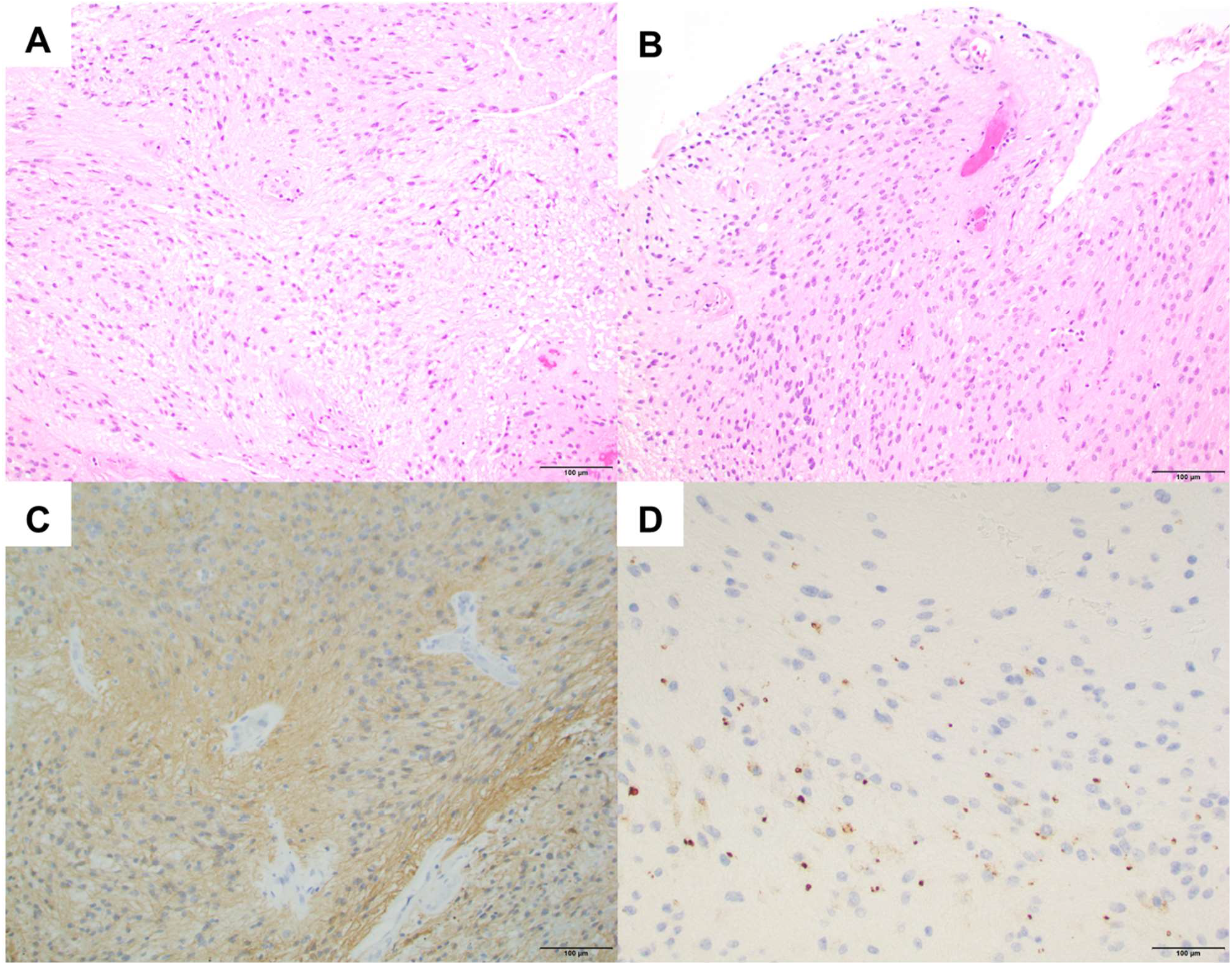
Histologic imaging of the SP-EPN sample, 20X magnification with scale. H&E staining showed compact glial neoplasm with fascicular architecture and fibrillar processes radially arranged around vessels (perivascular pseudorosettes) (A, B). GFAP immunostaining emphasizing radial arrangement of processes around vessels (C). EMA staining highlights microlumina in a dot-like pattern in well differentiated cells (D).

**Fig. 2.**
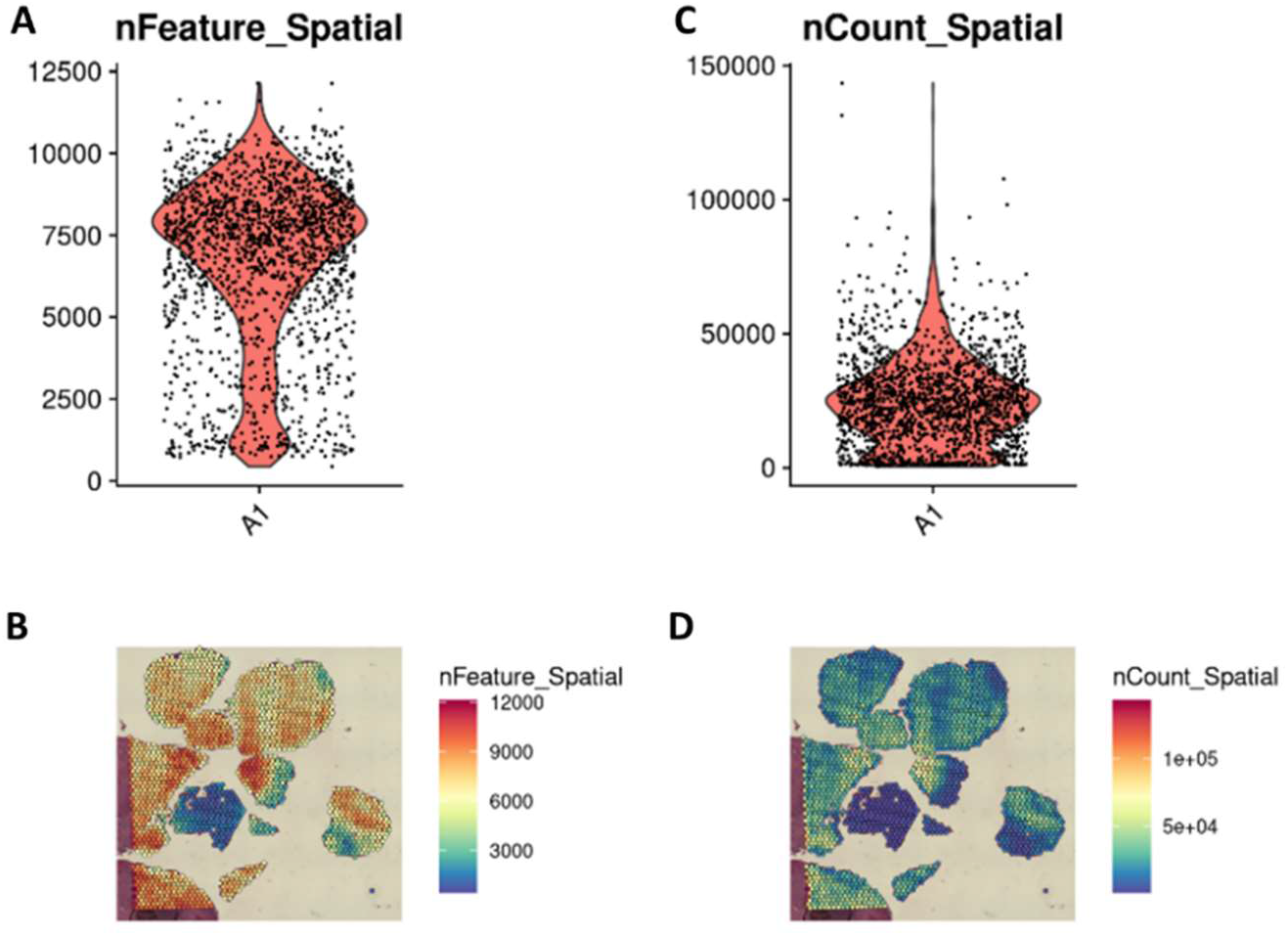
Plots assessing the quality of the SP-EPN sample. nFeature_spatial plots show the number of genes expressed in each spot of the capture area (A,B). nCount_Spatial reveals moderate level of reads throughout the sample (C, D).

**Fig. 3.**
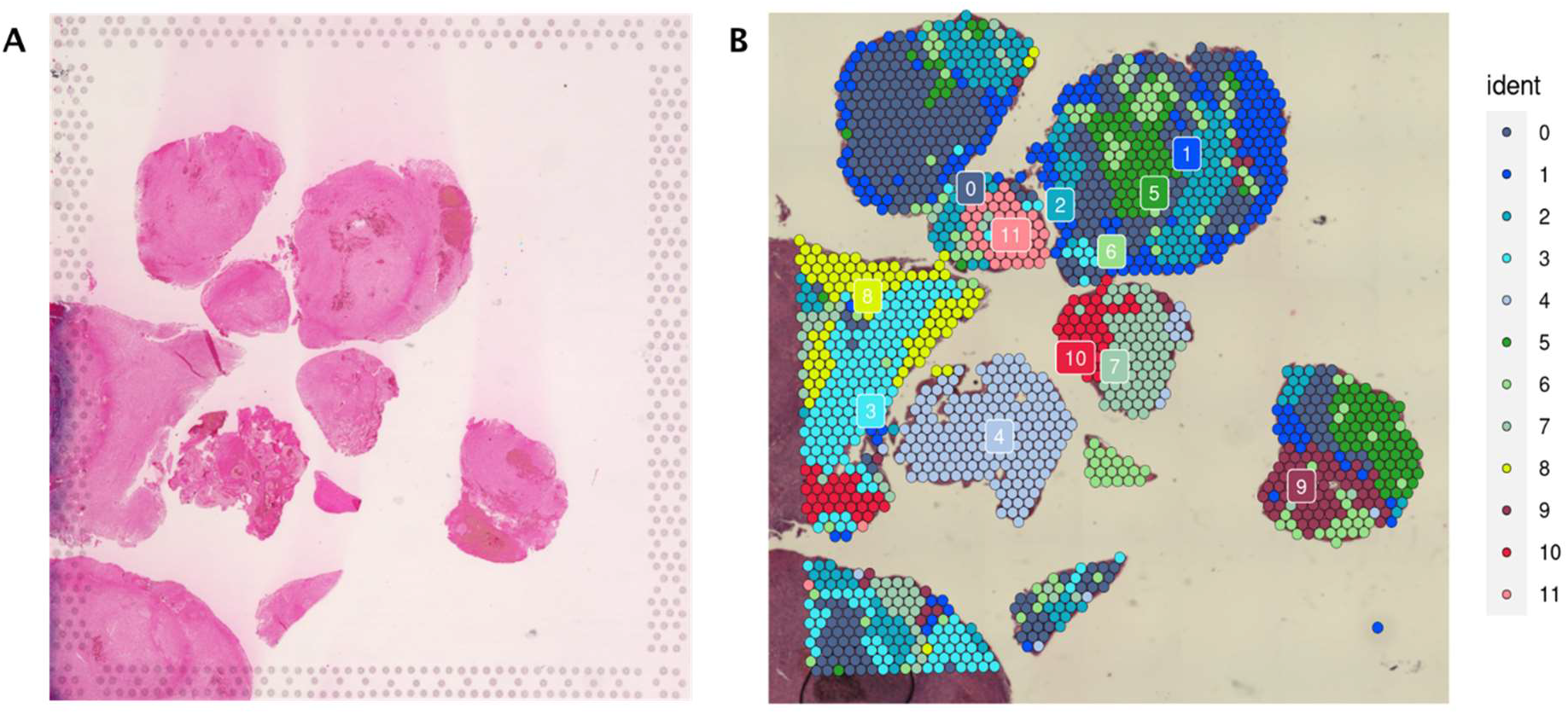
Cellular heterogeneity in SP-EPN. Histological examination of spinal ependymoma shows a fairly homogeneous tumor. However, spatial transcriptomics reveals marked heterogeneity with molecularly distinct cell groups.

To dissect the heterogeneity of the tumor further, we interrogated the ST data using specific gene markers of neural cell types found in the developing CNS. Specifically, we examined molecular markers of NECs (*SOX1, DACH1, ZBTB16*), RGCs (*FABP7, GFAP, SLC1A3*), EPCs (*PIFO, FOXJ1, CFAP157*), astrocytes (*SOX9, SLC1A3, GFAP*) and neurons (*MNX1, CHAT*) (Fig. 4). Genes related to neuroepithelial cells (*SOX1, ZBTB16, DACH1*) generally had low expression in the tumor. Interestingly, the RGC marker *FABP7* was found highly expressed in pockets of the tumor, often in central areas of the sample compared to outer region of the tumor sample. Some of these areas of *FABP7* expression allied with the RGC to astrocyte lineage marker, *SLC1A3*. Interestingly, *SLC1A3* also overlapped with markers of cells transitioning to the EPC lineage (*PIFO*). More mature astrocytic markers, such as *SOX9* and EPC markers (*CFAP157, FOXJ1*) showed diffuse expression within the tumor with stronger localization in areas adjacent to the pockets with neural stem cell and RGC marker expression. In contrast, neuronal markers were not highly expressed (Fig. 4).

**Fig. 4.**
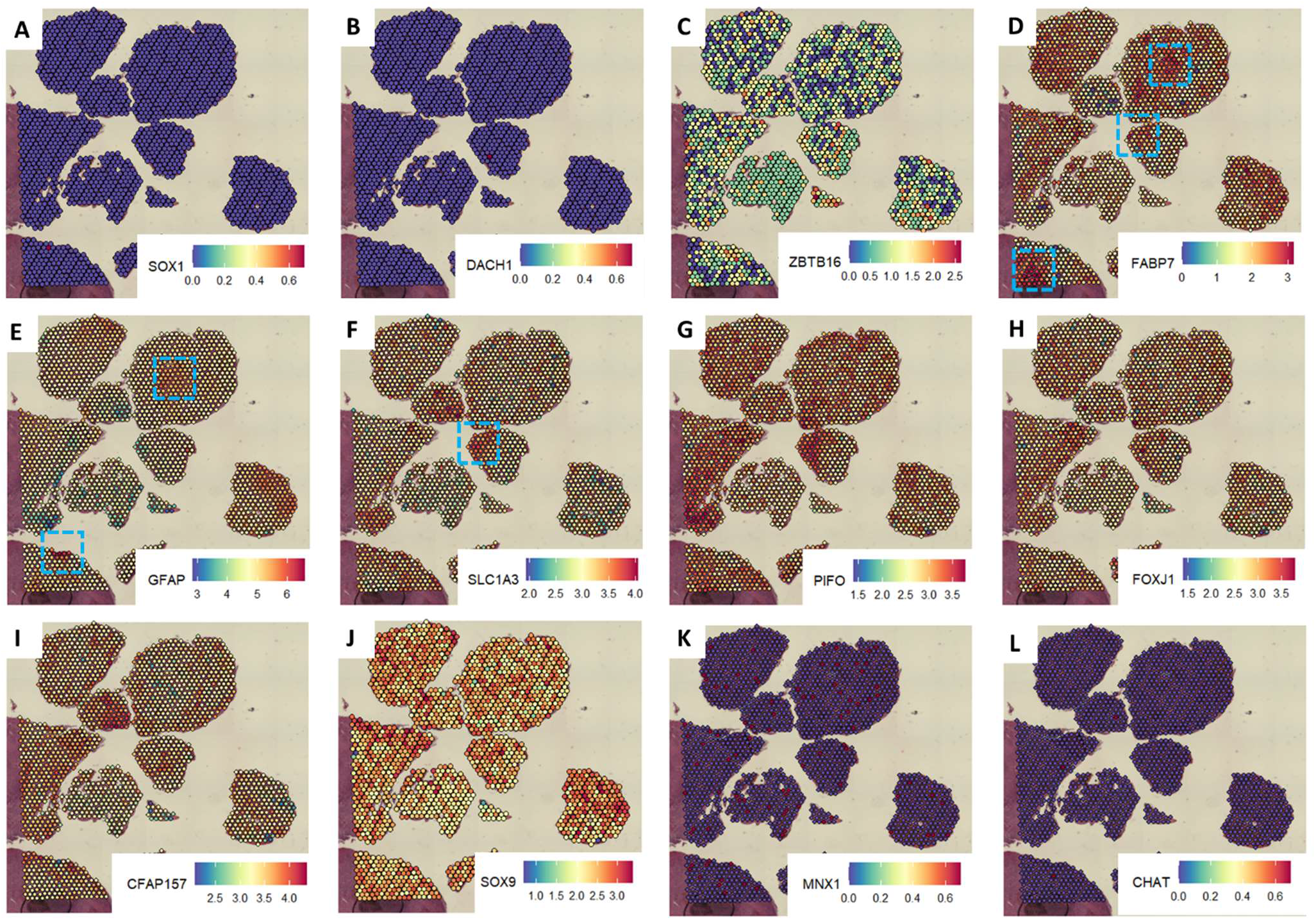
Spatial representation of individual neural and glial cell types in SP-EPN. Neuroepithelial cell markers SOX1, DACH1 and ZBTB16 are not highly expressed in the tumor (A,B). The radial glia markers FABP7, GFAP and SLC1A3 are seen highly expressed in smaller pockets within the tumor (D, E, F), blue brackets. More mature ependymal markers (PIFO, FOXJ1, CFAP157) and astrocyte markers (SOX9) are seen adjacent to these pockets of radial glia cells, with some overlap (G, H, I, J). Neuronal markers are not highly expressed (K, L). (Scale – fold expression (red = high, yellow = intermediate, blue = low)

To better define the relationship between NEC genes and more mature EPCs and astrocyte markers, we overlayed the marker expression of EPCs with astrocyte markers, which showed intermediate-high expression throughout the entire sample (Fig. 5). Interestingly, we noted smaller pockets of cell clusters with individual radial glia stem cell markers (*GFAP, Nestin, SOX2, SLC1A3 and FABP7*) (Fig. 5). Together, these findings suggest cellular heterogeneity within SP-EPN with most of the tumor displaying EPC and astrocytic markers, and smaller pockets with high RGC or neural stem cell marker expression.

**Fig. 5.**
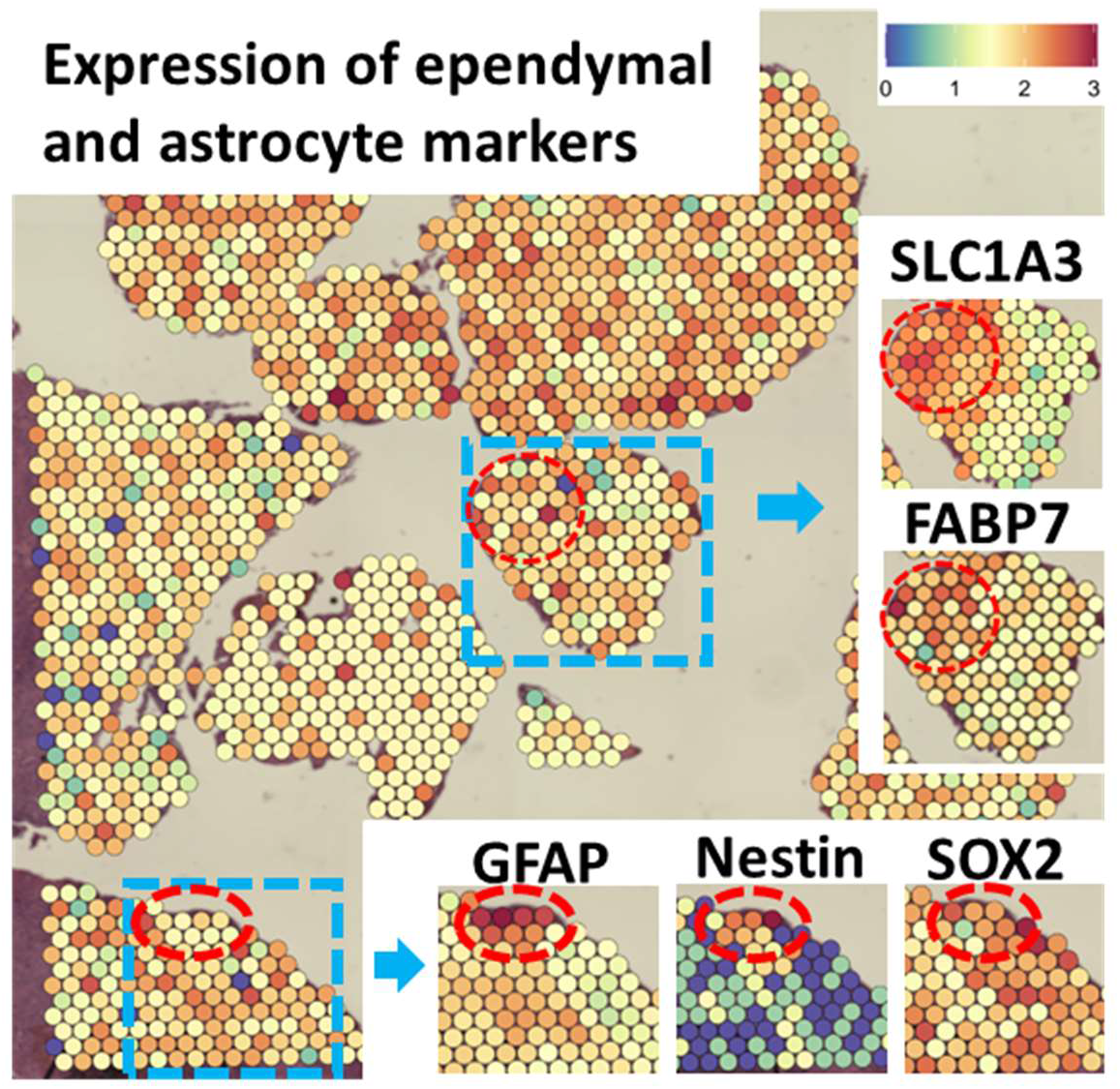
Spatial transcriptomics of spinal ependymoma (SP-EPN) from a NrS patient shows RG markers GFAP, Nestin, SOX2, SLC1A3 and FABP7 spatially represented in a smaller clusters of tumor cells. Each dot is 55 micrometers in diameter. Scale – fold expression (red = high, yellow = intermediate, blue = low)

## Discussion

Although SP-EPN are histologically homogenous tumors, our spatial transcriptomic analysis demonstrates molecularly distinct cell clusters with the tumor in a patient with NrS. Further, our study shows: 1) high expression of astrocytic and EPC markers within the tumor 2) smaller cell pockets with RGC or stem cell markers 3) relative low expression of neuronal markers. These findings suggest SP-EPN to predominantly be a disease of gliogenesis. Further, it points to the presence of a developmental hierarchy stemming from RGC-like cells in the EPC and astrocyte lineages.

In support of this, Garcia and Gutmann previously isolated neural progenitors from spinal cord of E12.5 *Nf2*^*flox/flox*^ mice.^16^ Interestingly, *Nf2* gene inactivation after injection of adenovirus containing Cre *ex vivo* led to increased growth of the cells and gliogenesis. These findings suggest that NF2 loss may lead to the development of aberrant glial progenitors with tumorigenic potential. However, the specific cell populations that are vulnerable to *Nf2* in this mouse model were not fully explored, and to date, a suitable mouse model for spinal ependymoma in NrS does not exist. We found expression of RGC, EPC and astrocytic genes in SP-EPN with *NF2* mutation, which is consistent with these previous findings. Importantly, our spatial maps show that these genes were expressed in spatially distinct regions. Astrocytic and EPC genes were generally found adjacent to smaller clusters with high RGC marker expression (*FABP7* and *GFAP*), with some overlap. The RGC to astrocyte lineage marker *SLC1A3* expression also coincided with some of the EPC genes, suggesting that there may be areas of overlap. Given that the capture area for each barcode is 55 micrometers, there may be both RGCs and EPCs within these areas, but there may also be cells that are transitioning from an RGC to EPC-like phenotype. In the developing mouse brain, EPCs are thought to arise from common embryonic RGCs.^17^ Our spatial transcriptomic data suggests a similar hierarchical organization may exist in ependymoma. Further studies with single cell RNA sequencing would be necessary to define the cellular hierarchy at single cell level.

### Limitations

We acknowledge our study has limitations. Firstly, the spatial transcriptomics platform we used did not provide single cell resolution and so rare population of cells may still be clouded by other cells within the cluster. Further, the trends we are see in this tumor may vary in other molecular subgroups of SP-EPN. We examined SP-EPN from a single NrS patient, further studies are warranted in other NrS and spontaneous SP-EPN sub-types. Furthermore, there may be sampling bias in the tumor based where the biopsy was taken. Nevertheless, our preliminary data highlights the importance of spatial heterogeneity within CNS tumors and beseeches caution when interpreting bulkRNA seq data for tumor origin. It provides a single proof of concept for a cellular hierarchy within spinal ependymoma and the presence of rare radial glia-like cells that requires deeper analysis in retrospective and prospective experiments. Further studies with single cell RNA sequencing methodology would be highly valuable.

### Conclusion

In conclusion, spatial transcriptomics performed on a SP-EPN tumor from a patient with NrS demonstrates marked cellular and molecular heterogeneity within the tumor. Our preliminary data suggests SP-EPN may be a disease of gliogenesis and that a developmental hierarchy may exist in the SP-EPN with RGC-like cells and more mature progenitors in the EPC lineage. Albeit a single proof of concept, these results offer important studies to pursue including single-cell analytics and animal studies targeting embryonic radial glia cells and the ependymal cell lineage.

## Methods

### Specimen Procurement

Spinal ependymoma sample was collected at our institution and formalin-fixed, paraffin embedded after surgery. Diagnosis of spinal ependymoma was decided through standard pathology processes based on 2016 World Health Organization CNS tumor classification criteria.^18^ Histological images (Fig. 1) were taken at 200X magnification.

### Sample Preparation

Samples were prepared as directed in the protocols of the 10X Genomics Visium CytAssist Spatial Gene Expression for FFPE (CG000495, CG000518, CG000520). SP-EPN sections were placed on a standard glass slide and H&E stained following deparaffinization. Afterwards, sections were coverslipped, imaged, and then decoverslipped, followed by decrosslinking. Human whole transcriptome probe panel was added to the deparaffinized, stained, and decrosslinked tissue. After probe hybridization and ligation, the glass slide with tissue sections was processed on the Visium CytAssist instrument to transfer analytes to a Visium CytAssist spatial gene expression slide with 11x11 mm capture areas. In the Visium CytAssist instrument, single-stranded probe ligation products were released from the tissue upon RNA digestion and tissue removal and were then captured on the Visium slide. Probes were extended by incorporating UMI, spatial barcode, and partial Illumina read 1. Libraries were generated from the extended probes, and then these libraries were quantified, and quality assessed with Qubit and Agilent TapeStation. The final libraries were sequenced on the Illumina NovaSeq 6000. 28-bp reads including spatial barcode and UMI sequences and 50-bp probe reads were generated with Illumina NovaSeq 6000 at the Center for Medical Genomics at Indiana University School of Medicine.

### Data Analysis

Space Ranger 2.1 was used to process the raw files following sequencing. Raw base sequence calls were demultiplexed to generate FASTQ files via bcl2fastq. Following this, the FASTQ files were used by spaceranger count to perform sequence alignment, tissue detection, fiducial detection, and spatial barcode/UMI counting. The reads were aligned to the human whole transcriptome probe set (Visium_Human_Transcriptome_Probe_Set_v2.0_GRCh38-2020-A.csv) and the human GRCh38 transcriptome. The reads aligned to the probe set were assigned to their respective genes and were associated with their specific spot based on their spatial barcode. Gene expression was quantified based on the number of UMIs detected within each spot.

The R package Seurat 4.9.9.9041 was used for data analysis following preprocessing with spaceranger.^19–22^ Sctransform was used to normalize the spatial transcriptome dataset. Clusters were identified using “FindClusters” and “FindNeighbors” functions.

## Acknowledgements

We want to thank the Center for Medical Genomics Core at Indiana University for their help and support with this study.

## Author Contributions

Conceptualization, N.B. and J.T.; Methodology, N.B., J.T., and H.G.; Software, N.B., J.W., H.G., and C.Z.; Investigation, N.B., J.W., H.G., R.B., C.Z. and J.T.; Writing – Original Draft, N.B., and J.T.; Writing – Review & Editing, N.B., J.W., H.G., R.B., C.Z., Y.L., W.C., and J.T.; Visualization, N.B., J.W., H.G., C.Z., and J.T.; Supervision, Y.L., W.C., and J.T.

